# Fast and accurate long-range phasing in a UK Biobank cohort

**DOI:** 10.1101/028282

**Authors:** Po-Ru Loh, Pier Francesco Palamara, Alkes L Price

## Abstract

Recent work has leveraged the extensive genotyping of the Icelandic population to perform long-range phasing (LRP), enabling accurate imputation and association analysis of rare variants in target samples typed on genotyping arrays. Here, we develop a fast and accurate LRP method, Eagle, that extends this paradigm to populations with much smaller proportions of genotyped samples by harnessing long (>4cM) identical-by-descent (IBD) tracts shared among distantly related individuals. We applied Eagle to *N*=150K samples (0.2% of the British population) from the UK Biobank, and we determined that it is 1–2 orders of magnitude faster than existing methods while achieving similar or better phasing accuracy (switch error rate ≈0.3%, corresponding to perfect phase in most 10Mb segments). We also observed that when used within an imputation pipeline, Eagle pre-phasing improved downstream imputation accuracy compared to pre-phasing in batches using existing methods (as necessary to achieve comparable computational cost).

---

Haplotype phasing is a fundamental question in human genetics^1^ and a key step in genotype imputation^2-5^. Most existing methods for statistical phasing apply hidden Markov models (HMM) to iteratively refine haplotype frequency models and improve phase calls^6-12^. This approach produces accurate phase inference at large sample sizes but is computationally challenging. “Long-range phasing” (LRP)^13^ is an alternative approach that harnesses long IBD tracts shared among related individuals; in such IBD regions, phase inference is straightforward and extremely accurate at sites for which at least one individual is homozygous. LRP has been successfully used in the Icelandic population to rapidly determine highly accurate phase and impute rare variants, producing insights into fine-scale recombination and enabling dozens of discoveries regarding numerous diseases^14-27^. However, because existing implementations of LRP rely on very long, easily identified IBD tracts (>10cM) in close relatives, LRP has previously only been successfully applied in isolated populations or populations with large fractions of individuals genotyped. In more general settings, existing LRP approaches are unable to phase a sizable fraction of sites^28^ and have been observed to achieve worse performance (both in terms of accuracy and run time) than conventional HMM-based approaches^29^.

Here, we develop a new algorithm, Eagle, that surmounts these challenges by combining the key ideas of LRP and conventional methods: Eagle begins with an LRP approach, making initial phase calls based on long (>4cM) tracts of IBD sharing in closely or distantly related individuals, and concludes with two approximate HMM decoding iterations to refine phase calls. We demonstrate the efficiency and accuracy of Eagle by phasing *N*=150K samples from the UK Biobank^30^ (see URLs); at large sample sizes, Eagle matches the accuracy of the best HMM-based methods and is far more computationally efficient (e.g., 14x faster than SHAPEIT2^12^). We also show that when phasing *N*=150K UK samples, Eagle imputes missing genotypes (in-sample) with accuracy *R*^2^>0.75 down to a minor allele frequency of 0.1%, and when used to pre-phase *N*=150K samples within a standard imputation pipeline, Eagle improves accuracy in downstream imputation (compared to pre-phasing using existing methods on batches of *N*=15K samples at comparable cost), with larger improvements expected as imputation reference panels grow. We have released Eagle as open source software (see URLs).

## Results

### Overview of methods

The basic idea of our approach is to harness IBD from distant relatedness (up to ≈12 generations from a common ancestor) that is pervasive within very large cohorts. IBD between a proband and other individuals provides a “surrogate family”^13^ for the proband, which can then immediately be used to call phase. While this approach is simple in principle, two major challenges have precluded its application to cohorts representing small fractions of large outbred populations. First, identifying IBD is difficult both in terms of accuracy and computational cost; moreover, the most widely used IBD inference methods rely on first phasing the data^31-33^. Second, LRP by itself can phase only sites at which the proband has at least one relative who is a homozygote; for cohorts representing a sizable fraction of a population, 2–5% of sites may be left unphased^13,15^, but for smaller cohorts, this fraction may exceed 25% even in isolated populations^28^, limiting the utility of LRP as a general-purpose method. Our algorithm, Eagle, overcomes the first challenge by employing a new, fast IBD-scanning strategy and overcomes the second challenge by introducing an approximate HMM computation that rapidly refines LRP phase calls.

The Eagle algorithm has three main steps (Figure 1). First, Eagle rapidly detects probable IBD tracts by identifying long regions of agreement at homozygous sites (i.e., identity by state, IBS≥1), scoring identified regions using allele frequency and linkage disequilibrium information, and checking overlapping regions for consistency; Eagle uses the detected IBD to perform accurate initial long-range phasing in high-IBD regions (Fig. 1a). Second, Eagle performs local phase refinement in overlapping ≈1cM windows by detecting complementary haplotype pairs (among haplotypes inferred in the previous step); specifically, for each diploid individual, Eagle re-phases the individual by searching for long IBD with haplotypes from step 1 and then checking for the existence of haplotypes complementary to the IBD hits (Fig. 1b). Third, Eagle finalizes phase calls by running two approximate HMM decoding iterations using up to 80 local reference haplotypes and aggressively pruning the search space to ≤200 states per position using a fast path-scoring scheme (Fig. 1c,d). All three steps are multithreaded and make use of bit operations to perform key computations in 64-SNP blocks. (For full details, see Online Methods and the Supplementary Note.)

**Figure 1:**
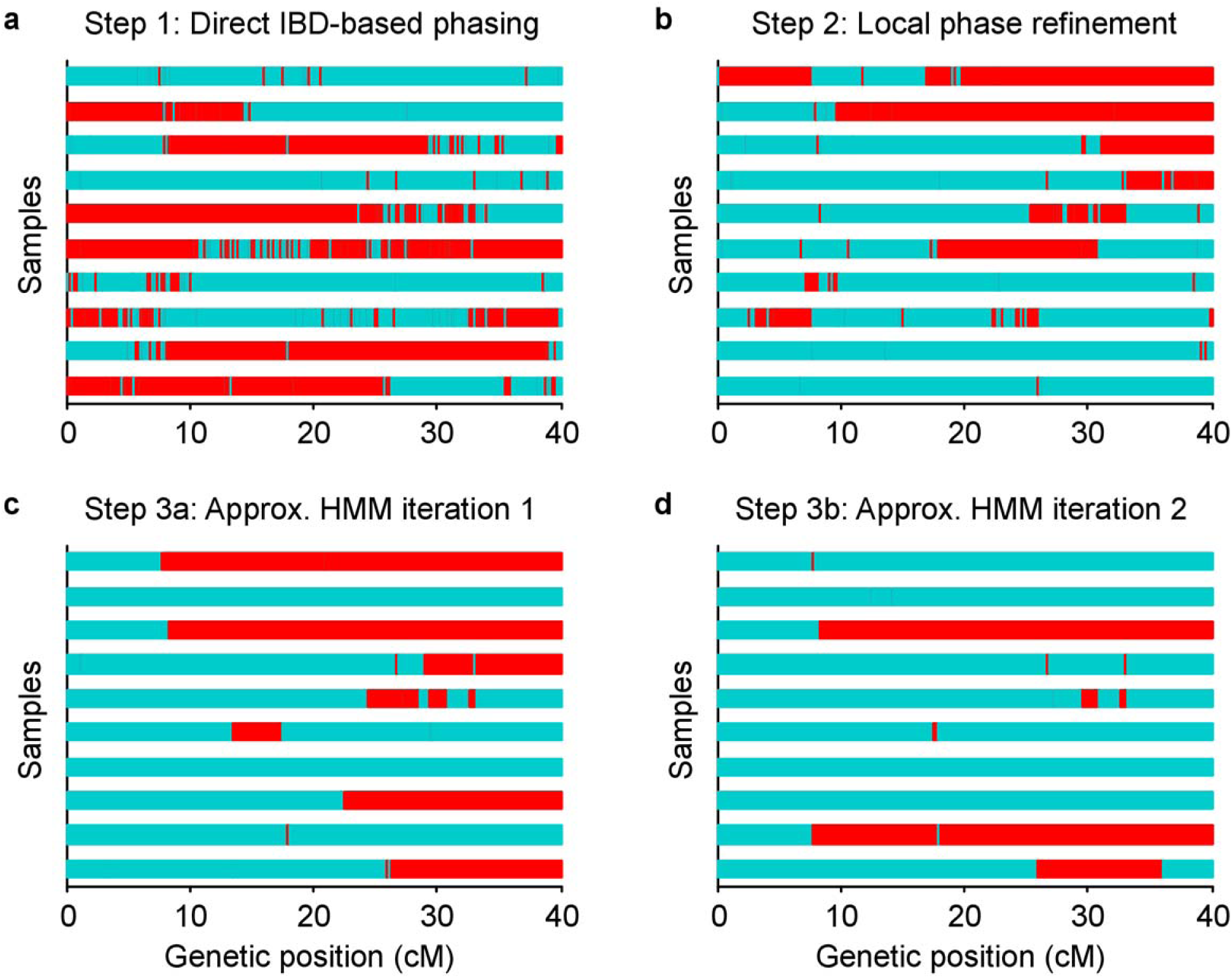
Eagle algorithm and example phase calls after each step. We show phase calls for ten trio children after each successive step of the Eagle algorithm (applied to phase the first 40cM of chromosome 10 in all *N*=150K UK Biobank samples except trio parents). At all trio-phased sites, red and blue indicate whether the first Eagle-phased haplotype for each child matches the maternal or paternal haplotype. (**a**) After the first step, a sizable proportion of each genome is covered by long segments of near-perfect phase; these segments are the regions in which long IBD is available from several relatives. (**b**) The second step, which uses both long and short IBD, fixes most of the phase switch errors in the first step. (**c,d**) The subsequent approximate HMM iterations further reduce the error rate.

### Computational cost

We benchmarked Eagle against state-of-the-art phasing methods—Beagle^8^, HAPI-UR^11^, and SHAPEIT2^12^ (see URLs)—on subsets of the UK Biobank data set containing *N*=15K, 50K, or 150K samples (Online Methods). After QC, this data set contained 627K autosomal markers with average heterozygosity 0.189 and minor allele frequency (MAF) distribution typical of genotyping arrays: 43K variants with MAF 0.1–1%, 235K variants with MAF 1–5%, and 349K variants with MAF 5–50%. (Our QC procedure excluded very rare variants with MAF<0.1%; see Online Methods.) For our first benchmark, we phased only the first 40cM of chromosome 10 (≈1% of the data, 5,824 SNPs spanning 18Mb) to allow as many methods as possible to complete in <2 weeks (using up to 10 cores on a single compute node; all methods except HAPI-UR support multithreading over 10 cores). We observed that Eagle achieved a 1–2 order of magnitude speedup over other methods across the sample size range (Fig. 2a and Supplementary Table 1), attaining a 14x speedup over SHAPEIT2 and a 12x speedup over HAPI-UR at *N*=150K. (Beagle was unable to phase 1% of the genome in 2 weeks at *N*=150K.) We note that (like other methods) Eagle has parameters that produce a trade-off in speed and accuracy (Online Methods); Eagle’s `–fast` mode achieved a further ≈2x speedup over the default while incurring only a slight loss of accuracy (Table 1). All methods exhibited superlinear but subquadratic scaling of running time with sample size, consistent with the presence of both linear and quadratic algorithmic components. (For a detailed discussion of the run time scaling of each of Eagle’s algorithmic steps, see Online Methods and Supplementary Table 2.) We also observed that Eagle achieved modest (2–8x) savings in memory cost compared to other methods (Fig. 2b and Supplementary Table 1). All methods exhibited memory cost scaling roughly linearly with sample size.

**Table 1:**
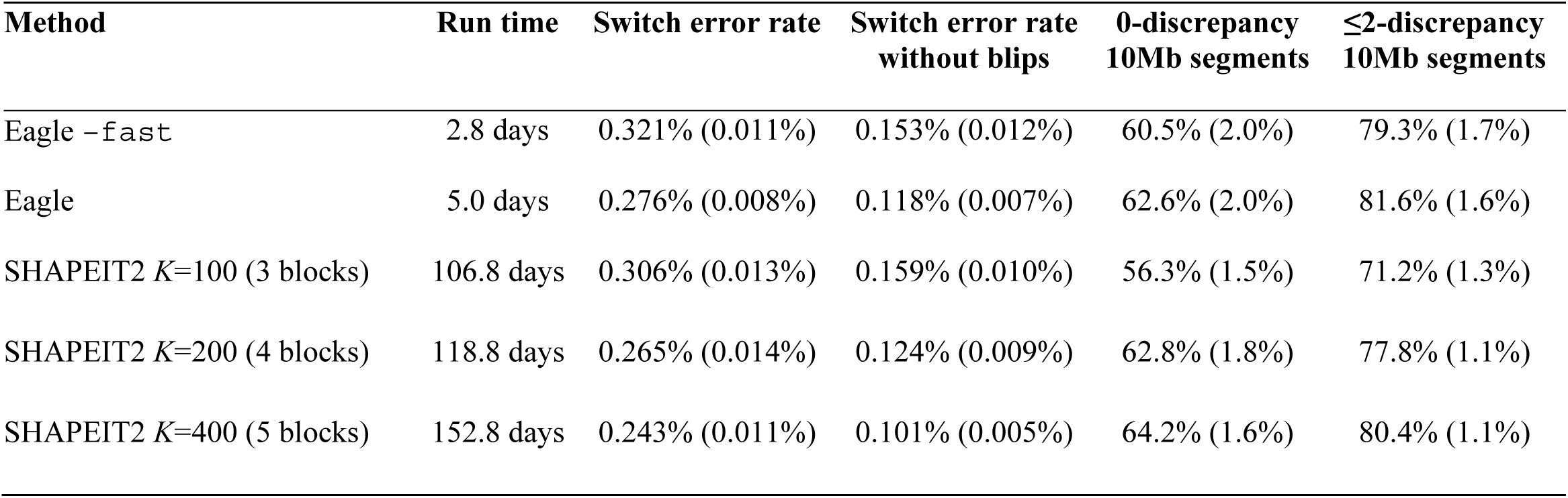
Computational cost and accuracy of Eagle and SHAPEIT2 on *N*=150K samples using various parameters.

**Figure 2:**
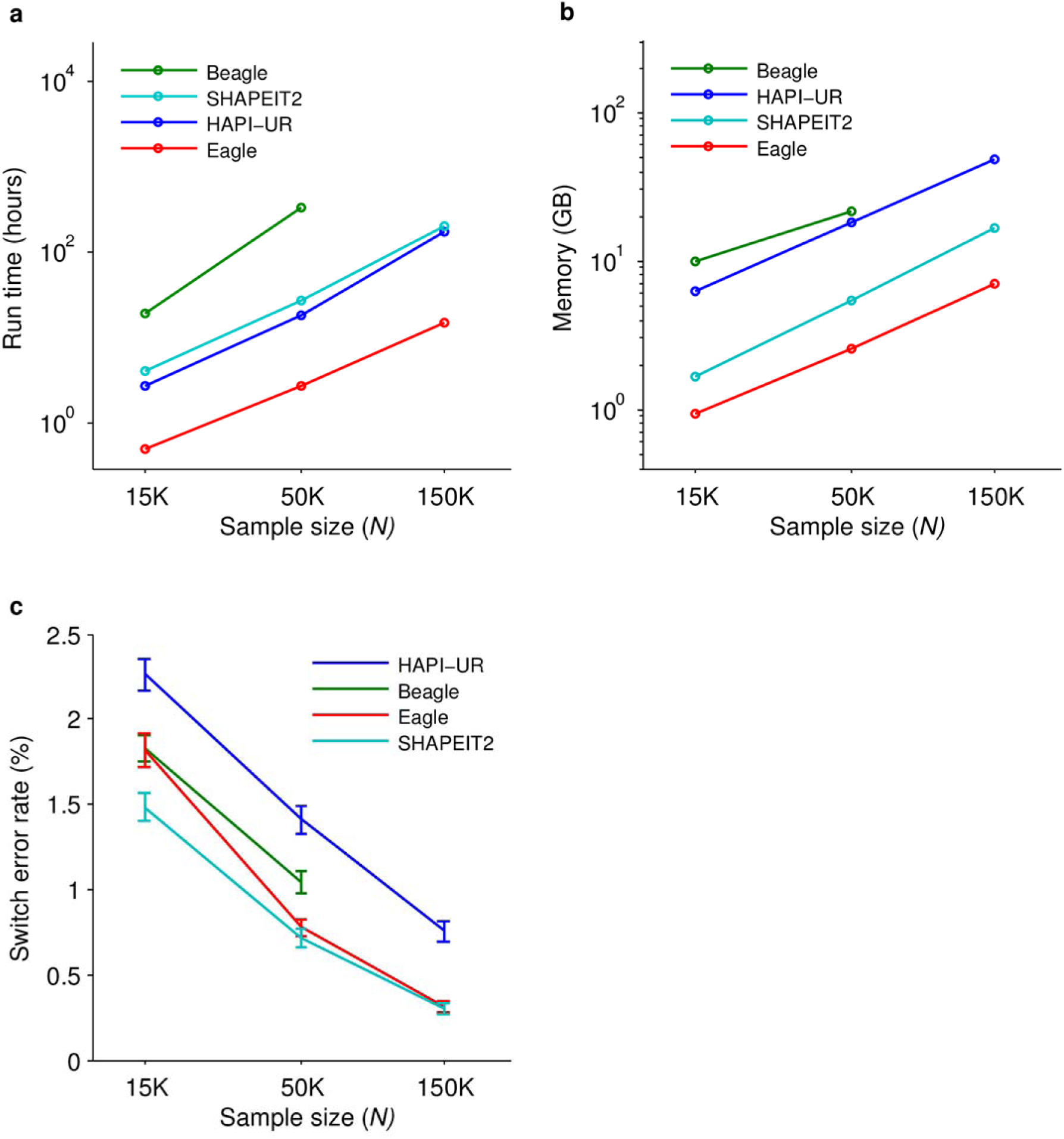
Computational cost and accuracy of phasing methods. Benchmarks of Eagle and existing phasing methods on *N*=15K, 50K, and 150K UK Biobank samples and *M*=5,824 SNPs on chromosome 10. (**a**) Run times and (**b**) memory using up to 10 cores of a 2.27 GHz Intel Xeon L5640 processor and up to two weeks of computation. (**c**) Mean switch error rate over 70 European-ancestry trios; error bars, s.e.m. All methods except HAPI-UR supported multithreading. As the HAPI-UR documentation suggested merging results from three independent runs with different random seeds, we parallelized these runs across three cores. (For the *N*=150K experiment, HAPI-UR encountered a failed assertion bug for some random seeds, so we needed to try six random seeds to find three working seeds. We did not count this extra work against HAPI-UR.) Numeric data are provided in Supplementary Table 1.

### Phasing accuracy

We assessed the accuracy of each phasing method using gold standard data from the 70 European-ancestry trios in the UK Biobank data set (all but one of which self-reported British ethnicity; see Online Methods). Specifically, we included all trio children and excluded all trio parents in each phasing run; we then assessed computational phase accuracy in trio children at all trio-phased sites (i.e., SNPs heterozygous in the child and homozygous in at least one parent, comprising ≈80% of heterozygous SNPs per trio child). We observed that when phasing *N*=150K samples over the same 1% of the genome as above, Eagle and SHAPEIT2 achieved near-identical, very low (≈0.3%) mean switch error rates (Fig. 2c and Supplementary Table 1). The accuracy of Eagle relative to SHAPEIT2 degraded slightly with decreasing sample size (as expected with limited IBD in an outbred population); interestingly, however, Eagle still achieved better accuracy than all methods except SHAPEIT2 at sample sizes of *N*=50K and *N*=15K, with only a 9% (s.e. 7%) increase in switch error rate relative to SHAPEIT2 at *N*=50K: 0.78% (0.05%) for Eagle vs. 0.71% (0.05%) for SHAPEIT2. To confirm these results, we performed a similar benchmark of Eagle and SHAPEIT2 on *N*=60K GERA samples of more diverse European ancestry^34,35^ (Online Methods) and observed similar results: 0.94% (0.04%) switch error rate for Eagle vs. 0.83% (0.03%) for SHAPEIT2, a 14% (2%) increase (Supplementary Table 3).

We next undertook a detailed comparison of phasing accuracy achieved by the two most accurate methods, Eagle and SHAPEIT2, when run on *N*=150K samples. For this comparison, we analyzed ten 10,000-SNP regions (of median length 44Mb) comprising 16% of the genome (Supplementary Table 4). To overcome the high computational cost of SHAPEIT2 *N*=150K analyses (Fig. 2a), we performed these benchmarks on the Lisa Genetic Cluster Computer (see URLs), which offered high-throughput parallel computation in batches of 16-core, 5-day jobs. Because SHAPEIT2 was unable to complete 10,000-SNP analyses within a single job, we applied a partition-ligation scheme to the SHAPEIT2 analyses, splitting each 10,000-SNP analysis into three overlapping blocks of 3,667 SNPs (with an overlap of 500 SNPs); we ligated the results using hapfuse v1.6.2 (see URLs). In these benchmarks, we observed that Eagle achieved a slightly lower switch error rate than SHAPEIT2 run with default parameters (*K*=100 conditioning states^12^): 0.276% (0.008%) for Eagle vs. 0.306% (0.013%) for SHAPEIT2 (*p*=0.03, one-sided paired *t*-test) (Table 1). One caveat regarding these comparisons is that applying partition-ligation to the SHAPEIT2 analyses may have incurred a slight loss of accuracy^12^; while computational limitations prevented us from running SHAPEIT2 on full 10,000-SNP regions, performing the full analyses in single computations could improve accuracy.

In light of the very low switch error rates achieved by both Eagle and SHAPEIT2 in *N*=150K analyses, we further investigated the nature of the errors made by each method. We observed that many of the switch errors accrued by Eagle were the result of “blips” involving one or occasionally two adjacent SNPs oppositely phased relative to surrounding SNPs (Fig. 1d); such errors can arise from genotyping error or from recent mutation or gene conversion. We therefore computed an alternative metric, “switch error rate without blips,” in which we ignored errors in which 1–2 SNPs were oppositely phased relative to ≥10 consistently phased SNPs on both sides. This assessment showed that 1–2 SNP blips (previously counted as two switches) accounted for the majority (≈60%) of Eagle’s switch errors; similarly, such errors accounted for roughly half of SHAPEIT2’s switch errors (Table 1). We further considered the metric “discrepancies within a 10Mb segment”^13^ and observed that both Eagle and SHAPEIT2 achieved perfect phase in the majority of 10Mb segments phased (Table 1 and Supplementary Table 5).

As both Eagle and SHAPEIT2 have parameters that trade off speed and accuracy, we also investigated the effect of running each method with non-default parameter settings. For Eagle, we benchmarked its `–fast` mode, which speeds up analysis by increasing the size of SNP blocks and limiting the approximate HMM computation (Online Methods). We observed that the `–fast` mode of Eagle completed analyses roughly twice as quickly as the default mode with a slightly higher switch error rate (0.321%, s.e. 0.011%) (Table 1). We also benchmarked slower parameter settings that decrease the size of SNP blocks or expand the approximate HMM computation; these modifications did not significantly improve accuracy (Supplementary Table 6). For SHAPEIT2, we increased the number of conditioning states from *K*=100 (the default) to *K*=200 or *K*=400, simultaneously increasing the number of partition-ligation blocks to 4 or 5 per 10,000-SNP region to keep per-job run times within the 5-day limit (Table 1). We observed that using *K*=200 conditioning states achieved accuracy similar to Eagle, while using *K*=400 states achieved the lowest switch error rate of all methods tested (0.243%, s.e. 0.011%; *p*=0.007 vs. Eagle, one-sided paired *t*-test) (Table 1). (We note that SHAPEIT2 *K*=400 analyses required ≈40% more computation time than default SHAPEIT2 *K*=100 analyses; while the run time scaling of SHAPEIT2 is asymptotically linear in *K* (ref.^12^), the quadratic component of the computation, which is independent of *K*, dominates at very large *N* and typical *K*.) We also considered increasing SHAPEIT2’s window size parameter from 2Mb to 4Mb, but results of a pilot experiment indicated that doing so substantially decreased accuracy (Supplementary Table 7).

Finally, we assessed the accuracy of analysis options for efficiently phasing *N*=150K samples. In addition to Eagle (run on all *N*=150K samples together, 1×150K), we considered batching approaches requiring up to 3x the running time of Eagle 1×150K. Based on our running time benchmarks (Fig. 2a and Supplementary Table 1), SHAPEIT2 or HAPI-UR analysis of the data in 10 batches of *N*=15K samples (10×15K) satisfied this constraint. We benchmarked each method on three chromosome-scale tests: the short arm of chromosome 1 (26,695 SNPs), chromosome 10 (31,090 SNPs), and chromosome 20 (16,367 SNPs), amounting to 12% of the genome. Our results (Supplementary Table 8) confirmed our previous benchmarks (Figure 2) and were consistent across chromosomes. In particular, we observed that Eagle analysis of all *N*=150K samples together completed 3x faster than SHAPEIT2 10×15K analysis while achieving a 77% (1%) decrease in switch error rate: 0.31% (0.01%) for Eagle 1×150K vs. 1.35% (0.05%) for SHAPEIT2 10×15K (Supplementary Table 8).

### Imputation accuracy

We next investigated the utility of Eagle for genotype imputation. To project the imputation accuracy that will be achievable in the UK population using LRP-based methods once a reference panel of *N*=150K sequenced UK samples becomes available (Supplementary Fig. 1), we performed in-sample imputation of masked genotypes in the UK Biobank data set (Online Methods and Supplementary Note). In these benchmarks (Supplementary Fig. 2 and Supplementary Tables 9–11), Eagle and SHAPEIT2 both achieved mean in-sample imputation *R*^2^>0.75 down to a minor allele frequency of 0.1%. As in our switch error benchmarks (Table 1), Eagle was slightly more accurate than SHAPEIT2 run with default parameters and achieved accuracy similar to SHAPEIT2 run with *K*=200 states; compared to SHAPEIT2 10×15K, Eagle 1×150K was much more accurate (Supplementary Fig. 2 and Supplementary Table 9). In-sample imputation on *N* samples bears some similarities to standard GWAS phasing and imputation on a target sample using a reference panel of size *N* (as both tasks entail copying shared haplotypes—identified based on data at typed SNPs—from a set of *N* samples); however, the two tasks also have several important differences and require different algorithms and software (Supplementary Fig. 1). We therefore caution that our in-sample imputation results may not be representative of GWAS imputation performance using an *N*=150K reference panel.

We also investigated the benefits of using Eagle for pre-phasing^5^ within an existing imputation pipeline: the Sanger Imputation Service, which currently supports imputation using up to *N*=32K sequenced reference individuals from the Haplotype Reference Consortium (HRC; see URLs). (We note that the HRC is predominantly European and contains a substantial fraction of UK samples but also contains samples of other ancestries; see URLs.) We considered two fast pre-phasing procedures: Eagle pre-phasing of all *N*=150K UK Biobank samples and SHAPEIT2 10×15K pre-phasing of *N*=150K samples. To benchmark imputation accuracy, we completely masked 700 SNPs (100 in each of seven MAF bins) in each of three chromosomes, pre-phased the remaining SNPs with Eagle and SHAPEIT2, imputed the same subset of *N*=15K pre-phased samples using the Sanger Imputation Service, and computed *R*^2^ between the masked SNPs and their imputed genotype dosages across curated British samples (Online Methods; see URLs). This benchmarking procedure is commonly used to assess the accuracy of phasing and imputation pipelines^5,9^. We observed that when imputation was performed using the largest reference panel available (the *N*=32K HRC), Eagle pre-phasing using all *N*=150K samples improved imputation *R*^2^ by increasing amounts for increasingly rare SNPs, with a gain of 0.020 (0.002) in *R*^2^ for MAF 0.1–0.2% SNPs (*R*^2^=0.594 (0.012) for Eagle 1×150K vs. *R*^2^=0.574 (0.012) for SHAPEIT2 10×15K; Supplementary Table 12). (We caution that these results are based on the preliminary release (r1) of the HRC; development of the HRC is still underway, and performance may improve in future releases.) When imputation was performed using only the *N*=4K UK10K reference panel (see URLs), gains were roughly half as large (Supplementary Table 13). Finally, to verify that similar improvements could be obtained at genome-wide SNPs (vs. the subsets of SNPs we masked), we ran the 1000 Genomes GBR samples through the same pipeline (after pre-phasing them together with the UK Biobank samples) and again observed a modest improvement using UK10K imputation (Supplementary Table 14). (We were unable to perform this experiment using HRC imputation because the HRC contains the 1000 Genomes data.) These results demonstrate that high-accuracy pre-phasing is already beneficial for GWAS imputation at current reference sizes (*N*=4K UK10K samples and *N*=32K diverse European HRC samples) and that gains will increase as reference panels grow, corroborating our in-sample imputation results projecting high future accuracy with *N*=150K reference samples.

## Discussion

We have developed a fast and accurate LRP-based phasing method, Eagle, and demonstrated that LRP can be effective in a cohort representing a small fraction of a large outbred population. Ever since Kong et al.^13^ established the efficacy of LRP in the Icelandic population—speculating that “having as little as 1% of a population genotyped may be adequate for the method to yield useful results”—the extension of LRP to more general settings has been eagerly anticipated but up to now unrealized^1^. We have successfully applied Eagle to phase 0.2% of the UK population and demonstrated its utility for enhancing the accuracy of downstream imputation. We note that LRP in 0.2% of the UK population cannot match the accuracy that was achieved in 11% of the Icelandic population^13,15^ (which further improved with genotyping of >30% of the Icelandic population^25^); however, these results give reason for optimism that LRP-based phasing accuracy in the UK (and other large outbred populations) will continue to improve as more individuals are genotyped.

Eagle is a very different method from the “pure” LRP approach of Kong et al.^13^: in order to create an algorithm that could harness limited, often distant relatedness, we needed to combine aspects of LRP and conventional HMM-based phasing, confirming the hypothesis that “IBD-based phasing can be extended…by using more sensitive methods for detecting IBD and combining IBD-based phasing with population haplotype frequency models”^1^. Indeed, these ideas have implicitly begun to converge within sophisticated HMM-based methods (e.g., SHAPEIT2), as has recently been observed^29^. SHAPEIT2 takes a “bottom-up” approach in which it steadily improves phase accuracy over the course of a few dozen MCMC sampling iterations, iteratively copying phase information from progressively more accurate sets of best reference haplotypes. This procedure eventually achieves high-accuracy phase for a proband’s (distant) relatives, selects them as reference haplotypes, and uses them to phase the proband^29^. In contrast, Eagle takes a “top-down” approach, first scanning all pairs of individuals for long IBD tracts and using them to phase long stretches of genome, and then applying only two iterations of approximate HMM decoding to correct errors and fill in unphased regions (Figure 1). Thus, at a high level, the key methodological contribution of Eagle’s “top-down” approach is its use of LRP to greatly improve speed (by over an order of magnitude) by eliminating the need to slowly build phase accuracy over many HMM sampling iterations. This speedup is essential at large sample sizes: due to computational constraints, the production phasing of UK Biobank samples was not performed using the most accurate method available, SHAPEIT2; instead, a new (currently unreleased) method was developed, SHAPEIT3, which was reported to achieve a higher switch error rate of ≈0.4% (see URLs). At very large sample sizes to come, experience from Iceland indicates that HMM iterations may not be necessary at all^13,15,25^; instead, optimizing accuracy will require solving problems of a different nature, e.g., resolving conflicts in IBD information.

Beyond our immediate goal of fast and accurate phasing, we envision that the primary downstream application of Eagle will be genotype imputation in the UK Biobank and future population cohorts of similar or larger size. We have demonstrated the utility of Eagle within current imputation pipelines and the promise of this approach for use in future data sets (e.g., imputation using *N*=150K reference samples). However, realizing this potential will require additional work. First, as currently implemented, Eagle is optimized for phasing array data and will need to be modified to phase sequence data. In particular, the method will need to be modified to incorporate additional information available from paired-end reads^36^ and from rare variants—which can greatly IBD-calling—while accounting for increased error rates. Simulations with increased genotyping error suggest that the Eagle algorithm is in principle quite robust to error (Supplementary Table 15), but additional tuning will undoubtedly be necessary. Second, an imputation algorithm capable of rapidly and accurately imputing pre-phased target samples using very large imputation reference panels will be needed. Several efforts to develop such methods are currently underway: the Sanger Imputation Service (see URLs) is already using a new (unpublished) imputation algorithm based on the Positional Burrows-Wheeler Transformation (PBWT)^37^—which like Eagle applies fast string matching algorithms in favor of exact statistical modeling—and the Beagle v4.1 imputation software^38^ and the Minimac3 imputation software (unpublished but in use by the Michigan Imputation Server; see URLs) likewise aim to satisfy these requirements. Finally, the sequence data itself will need to be generated. However, very large scale sequencing projects are already underway: e.g., Genomics England plans to sequence 100,000 genomes by 2017 (see URLs).

While Eagle provides new levels of efficiency (and accuracy compared to fast alternatives) for phasing very large cohorts, we note a few limitations. First, Eagle relies on the IBD present within very large data sets to achieve high accuracy; on smaller data sets (e.g., *N*=15K), we recommend SHAPEIT2, which provides higher accuracy and is computationally tractable for such data sets. Second, along similar lines, we observed that when phasing all *N*=150K UK Biobank samples together, Eagle achieved lower accuracy than SHAPEIT2 on the <10K samples of non-European ancestry (due to limited IBD). In practice, such samples are easily detected (e.g., by using FastPCA^35^ or SNPweights^39^) and could be phased separately with SHAPEIT2. Alternatively, a hybrid algorithm that uses the Eagle approach for most of the phasing computation but switches to the SHAPEIT2 model in segments of genome lacking IBD would be ideal; developing such an algorithm is a direction for future work. Finally, despite Eagle’s speed, its computational complexity contains a quadratic term (like all other published methods) and will become daunting for million-sample data sets. Most simply, this issue could be sidestepped by phasing very large samples in batches of a few hundred thousand samples at a time, but we expect that further algorithmic improvements will be possible, e.g., limiting the set of haplotypes considered as potential surrogate parents via clustering methods (as in SHAPEIT3; see URLs). Despite these limitations, we expect that Eagle in its current form—already much faster than existing methods with equal or better accuracy—will be a useful tool for large-sample phasing, and we believe further innovations will amplify the advantages of LRP-based phasing and imputation.

## URLs

Eagle v1.0 software and source code, http://www.hsph.harvard.edu/alkes-price/software/.

SHAPEIT v2 software, http://mathgen.stats.ox.ac.uk/genetics_software/shapeit/shapeit.html.

HAPI-UR v1.01 software, http://code.google.com/p/hapi-ur/.

Beagle v4.0 software, http://faculty.washington.edu/browning/beagle/beagle.html. hapfuse v1.6.2 software, http://bitbucket.org/wkretzsch/hapfuse/src.

PLINK2 software, http://www.cog-genomics.org/plink2.

SNPweights v2.0 software, http://www.hsph.harvard.edu/alkes-price/software/.

UK Biobank, http://www.ukbiobank.ac.uk/.

UK Biobank Genotyping and QC Documentation, http://www.ukbiobank.ac.uk/wp-content/uploads/2014/04/UKBiobank_genotyping_QC_documentation-web.pdf.

UK Biobank Phasing and Imputation Documentation (including brief description of SHAPEIT3), http://biobank.ctsu.ox.ac.uk/crystal/docs/impute_ukb_v1.pdf.

GERA data set, http://www.ncbi.nlm.nih.gov/projects/gap/cgi-bin/study.cgi?study_id=phs000674.v1.p1.

1000 Genomes data set, http://www.1000genomes.org/.

UK10K project, http://www.uk10k.org/.

Haplotype Reference Consortium, http://www.haplotype-reference-consortium.org/.

Sanger Imputation Service, http://imputation.sanger.ac.uk/.

Michigan Imputation Server, http://imputationserver.sph.umich.edu/.

100,000 Genomes Project, http://www.genomicsengland.co.uk/the-100000-genomes-project/.

Lisa Genetic Cluster Computer, http://geneticcluster.org/.

## Acknowledgments

We are grateful to G. Bhatia, S. Gusev, M. Lipson, B. Pasaniuc, N. Patterson, and N. Zaitlen for helpful discussions. This research was conducted using the UK Biobank Resource and was supported by US National Institutes of Health grants R01 HG006399 and R01 MH101244 and US National Institutes of Health fellowship F32 HG007805. Computational analyses were performed on the Orchestra High Performance Compute Cluster at Harvard Medical School, which is partially supported by grant NCRR 1S10RR028832-01, and on the Lisa Genetic Cluster Computer (http://www.geneticcluster.org) hosted by SURFsara and financially supported by the Netherlands Scientific Organization (NWO 480-05-003 PI: Posthuma) along with a supplement from the Dutch Brain Foundation and the VU University Amsterdam.

## Online Methods

### Eagle algorithm

We outline the three main steps of the Eagle algorithm here; full details are provided in the Supplementary Note. The first and second step each iterate through all individuals in the data exactly once, updating each individual’s phase in turn; the third step performs two such iterations. To help guide intuition, Figure 1 provides a snapshot of the progress of the algorithm after each step for our first *N*=150K phasing benchmark (Figure 2).

### Step 1: Direct IBD-based phasing using long IBD

For each proband in turn, Eagle scans all other (diploid) individuals for long genomic segments (>4cM) in which one (haploid) chromosome is likely to be shared IBD with the proband. Eagle then analyzes these probable IBD matches for consistency, identifies a consistent subset, and uses this subset to make phase calls. In our *N*=150K analyses, this step required ≈10% of the total computation time (Supplementary Table 2) and achieved near-perfect phasing within long swaths of genome covering most of each sample (corresponding to regions with IBD to several relatives) (Fig. 1a). In more detail, our algorithm applies the following four procedures to each proband in turn.

First, we run a fast *O*(*MN*)-time scan against all other individuals for long runs of diploid genotypes containing no opposite homozygotes (i.e., IBS>0). This filtering procedure is expedient for analyses of very large data sets as it operates directly on diploid data and thus requires little computation; a few variations of the approach have previously been developed^40,41^. Our implementation achieves a very low constant factor in its running time by using bit operations to analyze blocks of 16–64 SNPs simultaneously and using dynamic programming to record the longest ten IBS>0 stretches starting at each SNP block. We partition SNPs into blocks as follows: moving sequentially across the genome, we initialize each new block to contain the next 16 SNPs. We then continue to add subsequent SNPs to the block until it either contains 64 SNPs or reaches a maximum span of 0.3cM; upon reaching either limit, we end the current block and begin the next block.

Second, we compute an approximate likelihood ratio score for each potential IBD match identified by the above scan. This procedure is similar in spirit to Parente2^42^, which likewise computes approximate likelihood ratio scores to increase sensitivity and specificity of IBD calls. Our approach prioritizes speed over accuracy; instead of using a haplotype frequency model as in Parente2, we use only allele frequencies and LD Scores^43^ to compute an approximate likelihood ratio for the observed match having occurred due to IBD versus by chance. We apply this procedure within a seed-and-extend framework in which we begin with long IBS>0 matches but consider extending them beyond IBS=0 sites (to tolerate genotyping errors). We record all extended matches with length >4cM and likelihood ratio >10*N* (where *N* is the number of samples) as probable IBD matches.

Third, we analyze the set of identified probable IBD matches for consistency, truncating or eliminating matches until we reach a consistent set. For any pair of overlapping probable IBD matches between the proband and potential surrogate parents 1 and 2, the implied shared haplotypes can be (a) consistent with the proband sharing the same haplotype with both surrogates 1 and 2, (b) consistent with the proband sharing one of its haploytpes with surrogate 1 and other with surrogate 2, or (c) inconsistent with both of these possibilities. We first identify pairs of overlapping probable IBD matches in which scenario (c) occurs; for these pairs, we assume the longer match is correct and trim the shorter match until consistency under either scenario (a) or (b) is achieved. If any match drops below 3cM after during this trimming procedure, we discard the match. At the end of the procedure, all remaining pairs of trimmed matches are consistent. We then perform a final check for global consistency of implied phase orientations among all matches, i.e., we reduce (if necessary) to a subset of matches that can each be assigned to either a surrogate maternal haplotype or a surrogate paternal haplotype in a manner that respects pairwise constraints (a) and (b).

Fourth, we use the surrogate maternal and paternal haplotypic assignments of probable IBD regions to make phase calls. Whenever at least one surrogate is homozygous at a proband het, we use that surrogate to phase the site. If all surrogates are also heterozygous, we make a probabilistic phase call based on the allele frequency of the SNP and the difference between the numbers of (heterozygous) surrogate maternal haplotypes and surrogate paternal haplotypes.

### Step 2: Local phase refinement using long and short IBD

For each diploid proband in turn, Eagle analyzes overlapping ≈1cM windows of genome, searching for pairs of haplotypes (from the output of step 1) that approximately sum to the diploid proband within the window. Eagle then makes phase calls according to the haplotype pairs that most closely match the proband. In our *N*=150K analyses, this step required ≈20% of the total computation time (Supplementary Table 2) and reduced the switch error rate to ≈1.5% (Fig. 1b). In more detail, our algorithm applies the following three procedures to each proband in turn.

First, we run a fast *O*(*MN*)-time scan to find probable IBD with other haploid chromosomes (according to phase calls made in step 1). This procedure begins analogously to the first component of step 1; again, we look for long segments of IBS>0 (now between the diploid proband and haploid potential surrogates), now allowing a single mismatch site (IBS=0) within runs. We then attempt to extend the identified seed matches and record the ten longest matches covering each SNP block (as defined above).

Second, for each window of three consecutive blocks (containing a total of up to 192 SNPs spanning up to 0.9cM), and for each of the ten longest haplotype matches covering that window, we search for haplotypes approximately complementary (within the window) to the long haplotype. The idea is that often, only one of the proband’s haplotypes belongs to a long IBD tract; however, in such cases, the other haplotype is often shared in a short IBD tract, allowing confident phase inference if the complementary haplotype can be found to exist. Looking for a complementary haplotype in an error-tolerant manner amounts to performing approximate nearest neighbor search in Hamming space; to do so, we apply locality-sensitive hashing (LSH)^44,45^. In brief, LSH overcomes the “curse of dimensionality” by building multiple hash tables (here, ten per window) using different random subsets of SNPs (here, up to 32); then, when searching for a complementary haplotype, chances are high that at least one hash table will not include any SNPs with errors, allowing the approximate match to be found.

Third, we select the lowest-error complementary haplotype pair in each window (i.e., block triplet) and use it to phase the block in the center of the window. This procedure is fairly straightforward, with the only subtleties being that at error SNPs (i.e., proband hets for which both surrogate haplotypes have the same allele), we defer to the surrogate with higher confidence (from step 1), and when transitioning from one block to the next, we choose the orientation of the next complementary haplotype pair that best continues the current surrogate maternal and paternal haplotypes.

### Step 3: Approximate HMM decoding

For each diploid proband in turn, Eagle identifies candidate surrogate parental haplotypes (from the output of step 2) for use within an HMM (similar to the Li-Stephens model^46^). Eagle then computes an approximate maximum likelihood path through the HMM using a modified Viterbi algorithm (aggressively pruning the state space to increase speed) and calls phase according to the HMM decoding. Finally, Eagle post-processes the phase calls to correct sporadic errors by explicitly taking into account haplotype frequencies and long IBD. Eagle runs two iterations of this entire procedure. In our *N*=150K analyses, this step required ≈70% of the total computation time (Supplementary Table 2) and reduced the switch error rate to ≈0.4% after the first HMM iteration and ≈0.3% after the second (Fig. 1c,d). In more detail, our algorithm applies the following three procedures to each proband in turn (in each HMM iteration).

First, we compile a set of reference haplotypes for the proband for each SNP block. This procedure begins analogously to the first component of step 2, identifying long haplotype matches using a fast *O*(*MN*) search within a seed-and-extend framework. To ensure that both maternal and paternal surrogates are represented among the reference haplotypes, we augment the set of long haplotype matches with complementary haplotypes found using LSH. In total, we store *K*≤80 reference haplotypes per block.

Second, we compute an approximate Viterbi decoding of a diploid HMM similar to the Li-Stephens model^46^ using the sets of local reference haplotypes found above. A path through the HMM consists of a sequence of state pairs (one maternal reference haplotype and one paternal reference haplotype) at each location; we score a path according to the number of transitions on the maternal side, the number of transitions on the paternal side, and the number (and types) of Mendel errors between the proband and surrogate parents. An exact Viterbi decoding of this HMM using dynamic programming requires *O*(*MK*^3^) time (for *K*^2^ state pairs and *O*(*K*) possible transitions per position), which is too expensive for us; instead, we perform the dynamic programming within a beam search, pruning the search space from *K*^2^ state pairs to the top *P*=100–200 state pairs at each location and thus limiting the complexity to *O*(*MKP*). We then phase the proband according to the approximate Viterbi path.

Third, we post-process the phase calls to correct sporadic errors. Within each window of three consecutive blocks, we use LSH to determine the frequencies of ≈1cM haplotypes that match the Viterbi-inferred maternal and paternal haplotypes up to at most two errors. In rare cases, the haplotype frequencies give strong evidence to flip the phase of one or two SNPs, in which case we override the Viterbi phase call. Finally, we also check the Viterbi-inferred maternal and paternal haplotypes for consistency with the longest previously-identified IBD segments; in rare cases when the Viterbi phasing requires a phase switch >1.5cM from either end of a probable IBD segment, we override the switch.

### Fast mode of Eagle algorithm

Many parameters of the Eagle algorithm can potentially be modified to trade off accuracy and speed. For simplicity, we created a single `–fast` mode that roughly doubles Eagle’s speed by increasing the maximum SNP block span from 0.3cM to 0.5cM and reducing the comprehensiveness of the second HMM iteration (by reducing its beam search width from *P*=200 to 100 and only re-phasing the samples processed in the first half of the first HMM iteration).

### Scaling of Eagle run time

Each of the three steps of the Eagle algorithm involves an all-pairs *O*(*MN*^2^) computation (*M* = number of SNPs, *N* = number of samples) followed by an additional computation; the latter computation is inexpensive for step 1 and scales close to linearly with *N* for steps 2 and 3 (Supplementary Table 2). Thus, the distribution of time spent per step changes slightly with sample size, but no specific step is asymptotically a bottleneck. Summing across the three steps, the all-pairs *O*(*MN*^2^) computation constitutes slightly over half of the total computational cost at *N*=150K (Supplementary Table 2).

All components of the Eagle algorithm have run time linear in the number of SNPs (with a small constant factor via bit operations). For genotype array data consisting mostly of common SNPs, linear scaling is optimal; however, for rare variant-dense data (e.g., sequence data), sublinear scaling should be possible, as rare variants have much lower information content than common variants. We note that this scaling could be achieved with some additional engineering, e.g., by applying Eagle to only a subset of common and low-frequency variants and incorporating compressed rare variants post-hoc (in a manner similar to imputation).

### UK Biobank data set

We analyzed data from the UK Biobank, consisting of 152,729 samples typed at ≈800K SNPs. Using PLINK2^47^) (see URLs), we removed 480 individuals marked for exclusion from genomic analyses based on missingness and heterozygosity filters, leaving 152,249 samples (see URLs, Genotyping and QC). We restricted the SNP set to autosomal, biallelic SNPs with MAF≥0.1% and missingness ≤5%, leaving 627K SNPs (26,695 on the short arm of chromosome 1, 31,090 on chromosome 10, and 16,367 on chromosome 20). We identified 72 trios based on IBS0<0.001, sex of parents, and age of trio members (see URLs, Genotyping and QC). Of the 72 trio children, 69 self-reported British ethnicity, one self-reported Indian ethnicity, and one self-reported Caribbean ethnicity. The remaining trio child did not self-report any ethnicity, but her parents self-reported Irish and “Any other white background” as their ethnicities. UK Biobank genotyping and QC analyses indicated that self-reported ethnicity aligned closely with genetic ancestry (see URLs); however, UK Biobank also curated a subset of 120,286 self-reported British samples recommended for GWAS.

### GERA data set

We analyzed GERA samples (see URLs; dbGaP study accession phs000674.v1.p1) typed on the GERA EUR chip^48^. The data contained 62,318 samples, of which we removed 961 with <90% European ancestry as determined by SNPweights v2.0 (ref.^39^). Among this subset of samples, we identified 197 trios from independent pedigrees according to relationships provided with the data release. We analyzed chromosome 10, which contained 32,741 SNPs.

### Phasing software versions and parameter settings

We tested the latest version of each method (as of August 2015) using its recommended parameter settings. For Eagle (v1.0), SHAPEIT v2 (r790), and Beagle (v4.0 r1399), no command line arguments were required beyond file paths and threading settings (10 computational threads). For HAPI-UR (v1.01), we set the maximum window size to 80 (as recommended based on genotyping density) and combined results from three parallel runs of the algorithm using different random seeds^11^.

### Evaluation of phasing performance

For our benchmark analyses of *N*=150K UK Biobank samples, we removed 144 trio parents and phased the remaining 152,105 samples. For our benchmarks on *N*=50K or 15K samples, we phased all 72 trio children along with 1/3 or 1/10 of the remaining non-trio parent samples (50,752 or 15,270 samples in total). We evaluated phasing accuracy in trio children by comparing computational phase calls to trio phase calls (ignoring SNPs with Mendel errors); trio phase was available at ≈80% of heterozygous SNPs. For each child, we computed switch error rate by dividing the number of phase mismatches at consecutive trio-phased SNPs by the total number of trio-phased heterozygous SNPs minus 1 (ref.^1^), i.e., ≈15% of all SNPs (varying slightly among samples). In our results, we report mean switch error rates over the 70 European-ancestry trio children (according to self-reported ethnicity; see above). We applied an analogous procedure for our GERA benchmarks (differing only in that we removed all known relatives of the trio children—as the data contained a few extended pedigrees—leaving 60,929 samples).

### Evaluation of in-sample imputation accuracy

In our in-sample imputation benchmarks, we used the same SNP and sample subsets described above, but we modified the genotype data by randomly masking 2% of all genotypes (increasing the missingness of each SNP by ≈0.02). We then phased the masked data, obtaining imputed genotypes at all masked SNPs in the phased output. For each SNP, we computed adjusted *R*^2^ between actual and imputed masked genotype values according to the formula

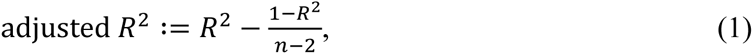

where *R*^2^ on the right is the usual coefficient of determination and *n* is the number of data points. (This adjustment corrects for upward bias due to finite sample size; for simplicity, we always use “*R*^2^” to refer to adjusted *R*^2^ elsewhere in this manuscript.) We computed means and standard errors of *R*^2^ over MAF strata, treating *R*^2^ from different SNPs as approximately independent given that the ≈2% subset of masked individuals varied from SNP to SNP. To assess in-sample imputation accuracy on a subset of samples (e.g., the 120K British samples curated by UK Biobank for GWAS), we computed *R*^2^ using only masked genotypes from samples in the subset.

### Evaluation of GWAS imputation accuracy

For computational efficiency, we performed all benchmarks of downstream imputation starting from a single data set, created as follows. First, we merged the 379 European-ancestry individuals from the 1000 Genomes Phase 1 integrated v3 release (see URLs) into the UK Biobank data set. Second, we entirely masked 700 random SNPs per chromosome, 100 in each of seven MAF bins (with MAF computed in the curated British samples). We phased all samples together using Eagle, and we phased a subset of *N*=15K samples (all 1000 Genomes samples plus 10% of the UK Biobank samples) using SHAPEIT2. Finally, we used the Sanger Imputation Service to impute the *N*=15K SHAPEIT2-phased samples and the same subset of Eagle-phased samples using both the UK10K panel (3,781 samples) and the Haplotype Reference Consortium (r1) panel (32,488 samples) with the PBWT imputation algorithm^37^ (see URLs). We assessed imputation *R*^2^ in *N*=12K curated British samples at the masked and imputed SNPs, computing means and standard errors across MAF strata as before (treating *R*^2^ from different SNPs as approximately independent given that each MAF bin contained <1 SNP per cM). We further assessed imputation *R*^2^ in UK10K-imputed 1000 Genomes GBR samples (*N*=89); since sequence data was available for these samples, we computed *R*^2^ at all UK10K-imputed SNPs in the 1000 Genomes data set. We computed means of *R*^2^ across MAF strata and estimated standard errors using a 100-block jackknife to account for linkage disequilibrium among SNPs.

